# Blackcaps (*Sylvia atricapilla*) with blood parasites have impaired oxidative status during autumn migration

**DOI:** 10.64898/2026.09.30.755661

**Authors:** Lucía Jiménez-Gallardo, Jimena López-Arrabé, Javier Pérez-Tris, Carolina Remacha

## Abstract

Parasites decrease host fitness, more so during physiologically challenging life stages such as migration. However, parasites are unevenly distributed among individuals, and hosts with different physiological condition may face different costs of migration with infections. We tested whether the relationship between infection by haemosporidian parasites and oxidative balance during autumn migration in blackcaps depends on individual age and sex, two major sources of variation in prevalence and physiological condition. We captured blackcaps at stopover in central Spain and measured infection status and physiological parameters. Blackcaps physiologically more prepared to depart from stopover had lower antioxidant capacity of plasma, probably because of physiological adjustments prior to the upcoming flight stage. *Plasmodium* parasites occurred only in coinfections, which were more frequent among young blackcaps. Regardless of individual sex and age, coinfected birds had lower antioxidant capacity of plasma, and blackcaps coinfected with *Plasmodium* had more oxidative damage to lipids in their red blood cells. Our results support the idea that haemosporidian infections impair the oxidative balance of blackcaps during autumn migration, which could impair individual performance and ultimately fitness, with increased population impacts during the first year of life when a larger proportion of individuals migrate coinfected with harmful *Plasmodium* parasites.

## 1. Introduction

Parasite infections impair host performance and ultimately fitness [1–3]. In birds, haemosporidian parasites (Apicomplexa) are a model for the analysis of the impact of parasites and its consequences on individual fitness [4]. Avian haemosporidians are vector-borne blood parasites of the genera *Plasmodium*, *Haemoproteus* and *Leucocytozoon* that invade the blood and other tissues of a wide range of bird species [5], impairing development, reproduction, and survival of their hosts [6–8]. Birds infected with haemosporidian parasites rely on various mechanisms to cope with infection [9,10], which require investing physiological resources that become unavailable for other vital functions, thereby entailing fitness costs [11,12].

During their life cycle, haemosporidian parasites activate the immune system of their hosts [13] and destroy red blood cells (RBC) and other invaded cells, inducing cell malfunction, anaemia, and lower haemoglobin concentration [14, 15]. These impacts are most important during the acute phase of infection, short after parasite invasion [2,8]. Parasite metabolism together with the immune response of the host produce reactive species (RS) [16,17], which could damage tissues if they accumulate in the organism [18]. Oxidative stress results from the imbalance between pro-oxidants and antioxidant defences, which can be endogenous or exogenous [19,20]. When confronting an oxidant situation, animals could either increase the production of endogenous antioxidants or promote the consumption of dietary antioxidants [19–21].

There are several stressful situations when birds may incur in oxidative damage such as development, senescence, reproduction or migration [22–24]. During migration, intense aerobic exercise and physiological adjustments during refuelling (including putting on fats prone to oxidation) make birds especially vulnerable to oxidative stress [25], which promotes up-regulation of endogenous defences or increased intake of dietary antioxidants during stopovers [26]. During those challenging moments, the impact of parasites is expected to be stronger, since limited antioxidant defences may strengthen the trade-off between immune defence and oxidative repair [27,28].

Few studies have analysed how parasites may alter oxidative balance in migrating birds [21], and whether this physiological link between infection and migration performance shows predictable individual variation [28]. When dealing with parasitic infections and oxidative stress, individual performance may depend on sex, age or body condition [8,19,29,30]. For instance, testosterone has an immunosuppressive effect on the humoral response [31], and females may have a stronger immune system than males [32,33], which can increase parasite impacts on males (as observed for *Plasmodium* parasitemia in mice [34]). Sexual differences in physiology can explain why males show higher haemosporidian prevalence than females in some bird populations (e.g. *Parus major* [35]). Host age is also important, as juveniles and adults differ in the degree of development of their immune system [36], their capacity to tackle oxidative stress [37–40], their experience in overcoming migration challenges [41], or how long they have harboured parasite infections [42]. Since younger birds would become infected for the first time, parasites might induce stronger innate (humoral) responses, with subsequent oxidative damage [18]. Moreover, migration might impose differential physiological costs to individuals depending on their sex and age combined, as adult males tend to arrive earlier to the breeding sites compared to females and juveniles [43], even shortening their refuelling time [44]. Then, males could be increasing migration costs to enhance their breeding success afterwards (e.g. [45]). Nevertheless, ontogenetic or sexual differences in physiological condition are far from universal among migrating birds (e.g. [46–48]. Taken together, the evidence indicates that it is important to consider individual sex and age to fully understand how birds cope with parasites and oxidative stress during migration.

We studied individual differences in the capacity of Eurasian blackcaps (*Sylvia atricapilla*) to maintain oxidative balance during autumn migration when infected by haemosporidian parasites. An earlier study in the same population, focused on dietary preferences of young male blackcaps, revealed lower endogenous antioxidants in infected individuals [21]. Here we analyse a larger sample of individuals to explore sex and age differences in the coping capacity of migrating blackcaps to maintain oxidative status. We distinguished between single infections and coinfections, considering the different parasite genera and lineages present in blood [49,50]. Coinfections might be important in a challenging context since they usually cause higher damage to their hosts [51]. We specifically investigated the infections of *Plasmodium* parasites, which are known to be more virulent than *Haemoproteus* and *Leucocytozoon* for various bird species [52,53] including blackcaps [21,54]. We predicted a negative relationship between parasite infection status and oxidative status measured by biomarkers of oxidative damage and antioxidant capacity. We expected a stronger effect for males due to the immunosuppressive effect of testosterone [31] and the better immune protection against parasites in females [33]. We also predicted a stronger effect for young blackcaps, since they are developing physiological and immunological mechanisms [36], and face infections for the first time which induces a stronger humoral response where RS production is expected to increase [18].

## 2. Materials and Methods

### 2.1. Field sampling

We conducted our sampling in a stopover site located in central Spain (40°20’08“N 3°24’27“W, 780 m a.s.l) during the peak of autumn migration of blackcaps [55] in October 2019 and 2020 (during 4 sampling days each year, 4-11 October in 2019 and 14-26 October in 2020). We captured individuals from approximately 9:30 until 15:30 (local time) with mist-nets, using blackcap song records to lure them. We captured 154 blackcaps in total, 89 in 2019 and 65 in 2020. Following [56], we determined the sex and age of individuals, differentiating between young birds born in the previous spring and older individuals (adults). Our sample included 66 females and 88 males; and 51 adults and 103 young birds. A sample of 49 males (48 of them juveniles) were kept in cages during ca. 30 min before processing, for a study on feeding preferences of young individuals [21]. The rest of birds were kept in cotton bags until processed. Different processing prior to sampling did not affect the physiological parameters of young males (ANOVA tests comparing caged and other young, all *p* ≥ 0.23). We weighed birds to the nearest 0.01 g and measured tarsus length with 0.01 mm precision. Finally, we took a blood sample (<1 % body mass) from the jugular vein. We used standard aluminium rings to individualise birds and then released them. We kept blood samples refrigerated in heparinised tubes until transported to the laboratory and processed during the same day. Samples were centrifuged to separate plasma from RBC and each subsample was stored at −80 °C until further analysis.

We used the residuals from the linear regression of log-transformed body mass on tarsus length (*r* = 0.18; *p* = 0.03) as a measure of size-corrected body condition of each individual [57]. For migrating blackcaps at stopovers, this measure of body condition is a proxy of the size of fat stores, which depends on time spent in the stopover site as blackcaps arrive lean and put on fat until they depart [58,59].

### 2.2. Haemosporidian parasite infections

We extracted total DNA from blood with SpeedTools Tissue DNA extraction kits (Biotools, Madrid, Spain) and diluted extractions to 25 ng/µl. Then, all the samples were screened for haemosporidian parasites following a multiplex PCR that distinguished between *Haemoproteus*, *Plasmodium,* and *Leucocytozoon* parasites [50], which increased the detectability of parasite coinfections of blackcaps [21]. All reactions included a negative control (ddH_2_O) and a DNA sample that was positive for all three genera of parasites. We run 4 µl of the PCR products on 2 % agarose gels stained with 100× GelRed^®^.

All samples found positive in the multiplex PCR were amplified with a nested PCR [60] that allowed us to determine the composition of each infection through sequencing of the MalAvi barcode, a fragment of 478 bp of the *Cyt-b* gene which is used to identify avian malaria lineages [49]. The final PCR products were sequenced using an Applied Biosystems™ 3730xl DNA analyser.

When several parasite lineages were detected either by the multiplex PCR or by mixed signal in sequencing electropherograms, we considered the sample as coinfected [50,61]. If only one parasite lineage could be confirmed, we considered individuals as single-infected. Accordingly, we classified the infection status of individuals using a factor with three levels: not infected, infected by one parasite, and coinfected by two or more parasites (only 5 individuals had 3 different parasites). Most infections involved *Haemoproteus* parasites, while *Leucocytozoon* was much less common. *Plasmodium* infections always occurred as coinfections alongside parasites from other genera (see results). Therefore, we conducted additional analyses focusing on *Plasmodium* coinfections, which are considered more virulent due to the synergistic effect of coinfection and the typically higher virulence of *Plasmodium* in birds in general [51–53], and in blackcaps in particular [21,54].

### 2.3. Oxidative status and physiological markers

All physiological analyses were carried out with exactly the same methods as reported in [21], and at the same time which allows direct comparison of the two studies. Prior to separating plasma from RBC, haemoglobin concentration was measured with Drabkin reaction (Spinreact, Girona, Spain), according to the manufacturer’s instructions. Haemoglobin levels could be related to parasite activity since haemosporidians induce anaemia [14], and they also vary during migration stopovers [62]. The assay carried out in duplicates showed high repeatability (*r_i_* = 0.89; *n* = 154; *p* < 0.001), with inter-assay CV = 12.61 % and intra-assay CV = 1.78 %.

Subsequently, the remaining blood was centrifuged for 10 min at 12 000 rpm to separate plasma from RBC. If haemolysis occurs during sample collection, the release of intracellular pro-oxidants and antioxidant molecules into the plasma may alter oxidative marker levels, potentially confounding the interpretation of the results [63]. Thus, the same observer visually determined the degree of haemolysis in plasma distinguishing three levels of redness (0: clear sample, 1: pale red stain, 2: red sample).

To standardize measurements performed on RBC (see below), protein content of RBC was quantified using the Bradford assay (Sigma-Aldrich, St. Louis, MO). Protein quantification measured in duplicates showed a repeatability of *r_i_* = 0.58 (*n* = 154; *p* < 0.001), with inter-assay CV = 17.21 % and intra-assay CV = 6.50 %.

Intracellular total glutathione level (tGSH) was measured as an endogenous antioxidant marker in RBC, following [63]. The tGSH levels measured in duplicates were standardised to the total protein content in the sample by dividing tGSH by protein values. The assay had a high repeatability (*r_i_* = 0.92; *n* = 153; *p* < 0.001), with inter-assay CV = 12.60 % and intra-assay CV = 8.31 %.

Lipid peroxidation was assessed as a measure of oxidative damage to cell membranes, through malondialdehyde (MDA) concentration. Blood parasites could induce peroxidation of RBC phospholipids due to the activation of the immune system and their own metabolism [64], which is particularly relevant during migration, when the integrity of cell membranes is key to the performance of the pectoral muscle [65]. In our population of migrating blackcaps, young males with higher MDA levels measured in plasma reinforce the intake of dietary antioxidants [21], supporting the value of MDA as a biomarker of oxidative status in this system. We quantified MDA in both plasma (plasma-MDA) and RBC (RBC-MDA), as indicators of lipid peroxidation and RBC membrane damage, respectively, following [39]. Samples were injected in duplicates, when possible (139 out of 149 samples of plasma-MDA, and 152 out of 154 samples of RBC-MDA). Repeatability of plasma-MDA was *r_i_* = 0.91 (*n* = 139; *p* < 0.001), with inter-assay CV = 27.44 % and intra-assay CV = 17.22 %. The repeatability of RBC-MDA was *r_i_* = 0.76 (*n* = 152, *p* < 0.001), with inter-assay CV = 20.58 % and intra-assay CV = 15.81 %. RBC-MDA values were standardized by total protein content.

Total antioxidant status (TAS) of plasma was quantified, reflecting circulating antioxidant defences such as vitamins C or E and carotenoids. In our system, infected young males have lower TAS values, supporting the idea that infections may deplete antioxidant defences [21]. TAS was measured following [66] with modifications by [67]. Samples, blank, and calibrator (Trolox) were run in duplicates (132 out of 148 samples). TAS measurements had high repeatability (*r_i_* = 0.96; *n* = 132; *p* < 0.001), with inter-assay CV = 14.19 % and intra-assay CV = 5.44 %.

Uric acid is an endogenous antioxidant and a product of the catabolism of amino acids in birds, usually positively correlated to TAS values [68–70], including in our study system [21]. Therefore, uric acid was measured in plasma to adjust TAS values using a commercial kit (Biosystems, Barcelona, Spain). Each sample was included in duplicates whenever possible (140 out of 151 samples). Uric acid measurements had high repeatability (*r_i_* = 0.95; *n* = 140, *p* < 0.001), with inter-assay CV = 4.07 % and intra-assay CV = 5.76 %.

Finally, plasma triglycerides were quantified as an indicator of the fattening status of the birds during stopover [71], using a commercial kit (Biosystems, Barcelona, Spain) according to manufacturer’s protocol. Given the relationship between triglycerides and MDA levels [63], triglycerides were considered in the analysis of this variable. Absorbance was measured in duplicates (138 out of 148) showing a high repeatability (*r_i_* = 0.98; *n* = 138; *p* < 0.001), with inter-assay CV = 1.72 % and intra-assay CV = 5.76 %.

### 2.4. Statistical analyses

We used the R package MASS v 7.3-60 [72] to fit an ordinal logistic regression model testing whether individuals of each sex or age class differed in their odds of bearing more parasites, using the multiple infection variable as a factor with three ordered levels, from not infected to coinfected. We used the Brant’s test implemented in the brant package v 0.3-0 [73] to check the proportional odds (parallel regression) assumption of the model. We fitted a logistic regression (with binomial error and logit link function) to analyse differences between sexes and age classes in the probability of harbouring *Plasmodium* infections.

The influence of haemolysis level and plate identity on each measurement was checked and controlled when needed. Since TAS and uric acid levels were strongly correlated (*r* = 0.78), the residuals of that relation were extracted as a corrected TAS measure. Although the correlation between plasma-MDA and triglycerides was weak (*r* = 0.15, F_1,142_ = 3.40, *p* = 0.07), plasma-MDA values were corrected using the residuals of their regression on triglycerides [63]. We checked the relation between all the variables that could be involved directly in the migratory metabolism of blackcaps, namely body condition (fat content), haemoglobin, uric acid, and triglycerides, and performed a Principal Component Analysis (PCA). The PC1 accounted for 34.0 % of variance in the correlation matrix (factor loadings: body condition = 0.61, haemoglobin = 0.63, uric acid = −0.58, and triglycerides = 0.50), and we interpreted PC1 scores as an index of individuals’ readiness to continue the migratory journey [74,75].

To assess if the effect of infections on the physiological condition of individuals depended on sex or age, we fitted lineal models (LM) with physiological parameters as dependent variables. We tested separately the effect of the multiple infection variable (not infected, single infected, or coinfected) and the effects of *Plasmodium* coinfections (present or absent), including age, year, and sex as covariates. We also included readiness to depart (PC1 scores) in the analyses where none of the variables included in this PCA was the dependent variable. For the complete assessment of the oxidative balance, when an oxidative damage biomarker was analysed as a dependent variable, all antioxidant biomarkers were included as covariates, and vice versa. We also included in the analysis the two-way interactions between infection and age, sex, and the physiological parameters included as covariates, when the corresponding interactions were significant. We tested which variables and interactions best predicted the response variable using the Akaike Information Criterion corrected for small sample size (AICc) [76] with the dredge function of the package MuMIn v 1.47.5 [77]. We averaged best models (ΔAICc ≤ 2) to compute significant effects [78], although none of the effects tested were considered significant if the best models included the null model. Whenever infection (the focal factor in our study) was included in any of the best models, we used the option “fixed” in MuMin to recalculate model-averaged estimates only with models including infection. All estimates were full averaged across models. Finally, when the multiple infection variable was retained in the model averaging step, we used the relevel option to estimate all pairwise between-level differences. All continuous variables were checked for normality, transformed when advisable (square root-transformed uric acid, tGSH, and TAS; log-transformed plasma-MDA and triglycerides), and z-standardised. Residuals of the models were distributed according to model assumptions. All statistical analyses were conducted using R 4.3.2 [79].

## 2. Results

Of the 154 blackcaps tested, 130 were infected with haemosporidian parasites (84.4 %), 43 harboured more than one parasite (33.1 % of the infected birds), and 22 were coinfected by *Plasmodium* parasites (16.9 % of the infected; see Supplementary Table 1 for further information of parasite lineages). Out of 103 young blackcaps tested, 91 (88.3 %) were infected, and 36.3 % of these harboured coinfections; while 39 (76.5 %) adults were infected, 25.6 % with coinfections. There were 10 (15.2 %) females not infected, 37 single-infected (56.1 %), and 19 coinfected (28.8 %); while males were 14 not infected (15.9 %), 50 single-infected (56.8 %), and 24 coinfected (27.3 %). For young birds, the odds of bearing higher levels of infection, i.e., being coinfected versus not infected, was 2.22 times that of adults (ordinal logistic regression estimate for young = 0.80; se = 0.35; 95 % CI [0.12, 1.50]; t_152_ = 2.27; *p* = 0.023; Fig. 1), holding constant the effect of sex, which was not significant (estimate for females = 0.24; se = 0.33; t_152_ = 0.73; *p* = 0.47). The probability of harbouring *Plasmodium* parasites was slightly higher for females than for males, although the difference was not significant (estimate for females = 0.86; se = 0.48; z = 1.80; *p* = 0.072). Controlling for sex in the logistic regression, both age classes showed similar probabilities of having *Plasmodium* infections (estimate for young = 0.51; se = 0.53; z = 0.97; *p* = 0.33).

**Figure 1.**
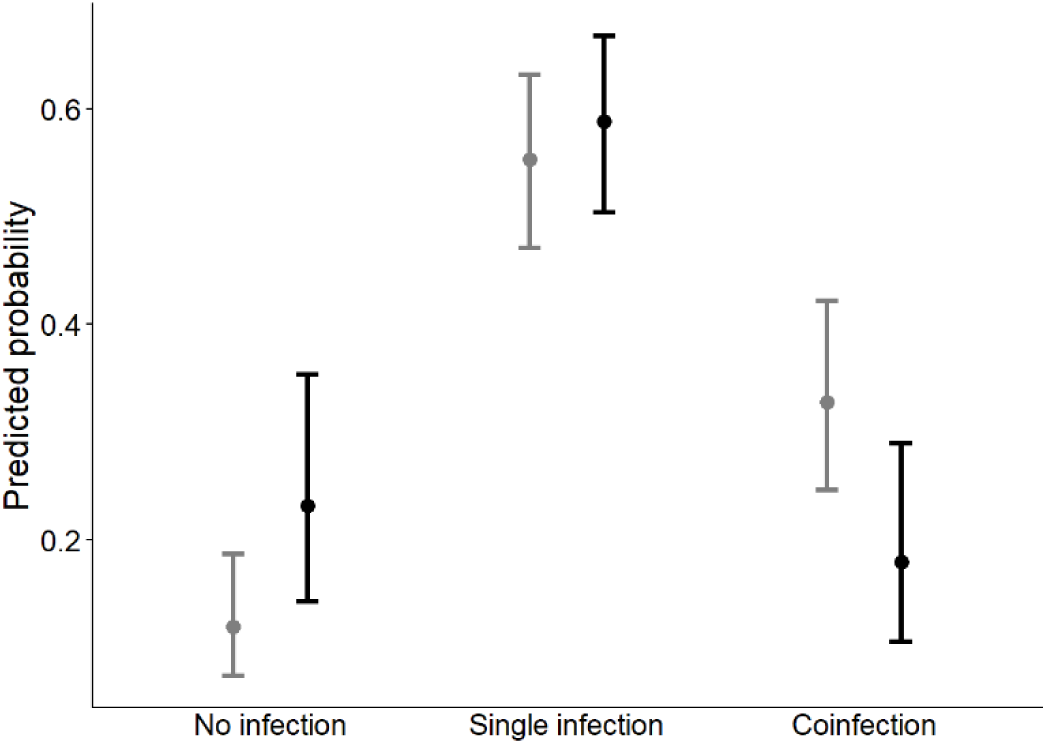
Estimated probability of bearing different levels of parasite infection (no infection, single infection, or coinfection) estimated for young (grey) and adult (black) blackcaps in an ordinal logistic regression model with age and sex (not significant) as predictors of infection status. Bars denote 95 % confidence intervals.

Models with ΔAIC ≤ 2 analysing variation in physiological parameters included the null model for haemoglobin, uric acid, plasma-MDA (in the case of total infections, but not in *Plasmodium* infections), and triglycerides (Supplementary Tables 2 and 3), and therefore were not further examined. Interactions between infection status and physiology, sex, or age were not significant and were not included in the final models (Supplementary Table 2 and 3). When testing for the different levels of the multiple infection parameter (fixed for model averaging), we found that birds infected with multiple parasites had lower TAS compared to not-infected individuals (Fig. 2A). Moreover, birds infected by *Plasmodium* parasites had higher values of plasma-MDA (Fig. 3). Finally, we found a positive relation between tGSH and RBC-MDA (*p* < 0.001; Supplementary Fig. S1 and S2), and birds closer to departure (i.e., with higher readiness PC1 scores) had lower levels of antioxidant capacity of plasma (TAS; Fig. 2A).

**Figure 2.**
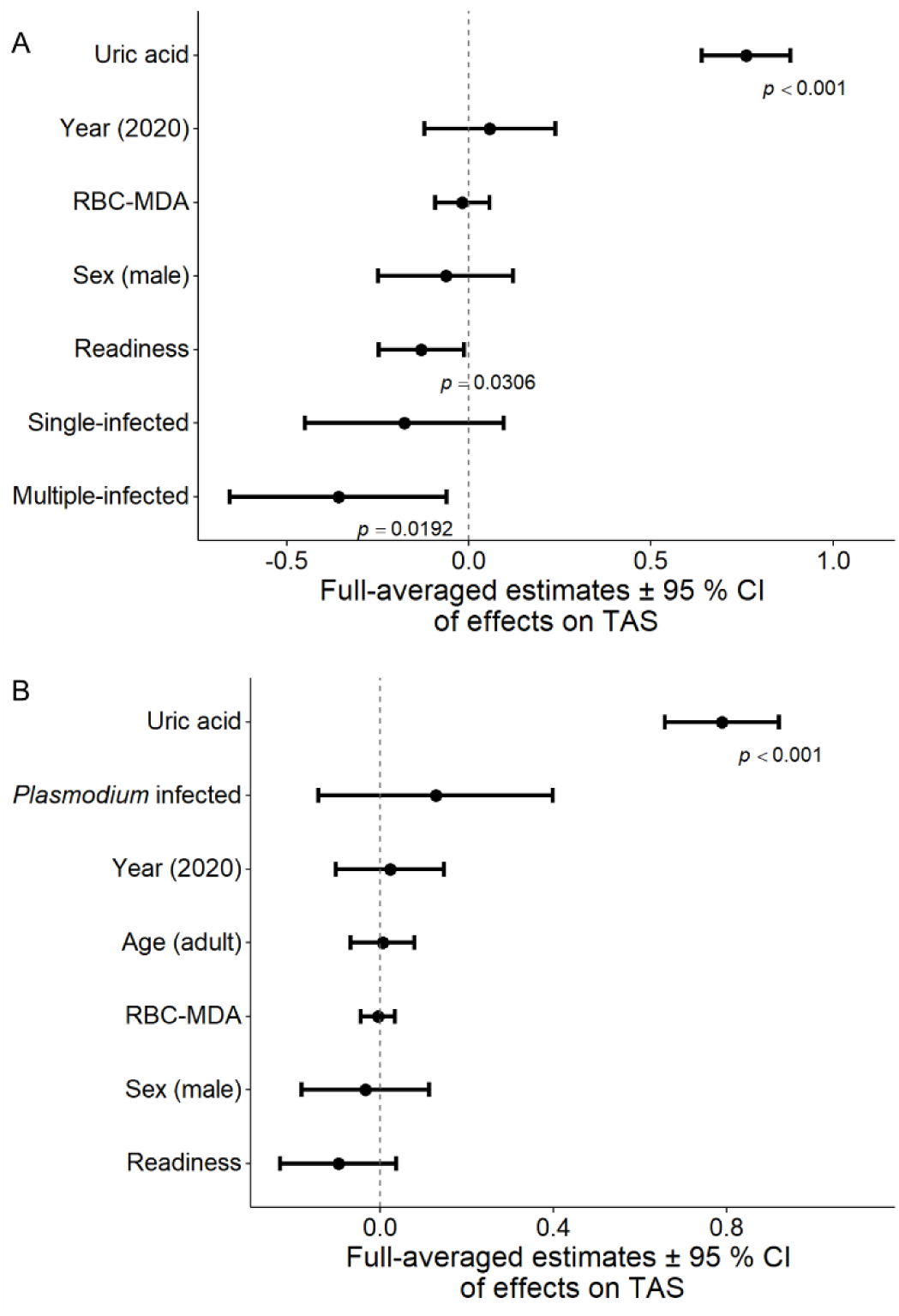
Standardized full model averaging estimates with 95 % confidence intervals of the effects of variables included in the best models for variation in TAS (square root transformed) of blackcaps, when these were tested using multiple infection status (A) or *Plasmodium* infection (B) as parasite-related predictors. The *p*-values of statistically significant effects are shown. The effect of multiple infection status is shown with not infected as the reference level. Changing the reference to single-infected: 0.18 ± 0.14, *p* = 0.21 not-infected; −0.18 ± 0.11, *p* = 0.11 multiple-infected. Uric acid was square root transformed.

**Figure 3.**
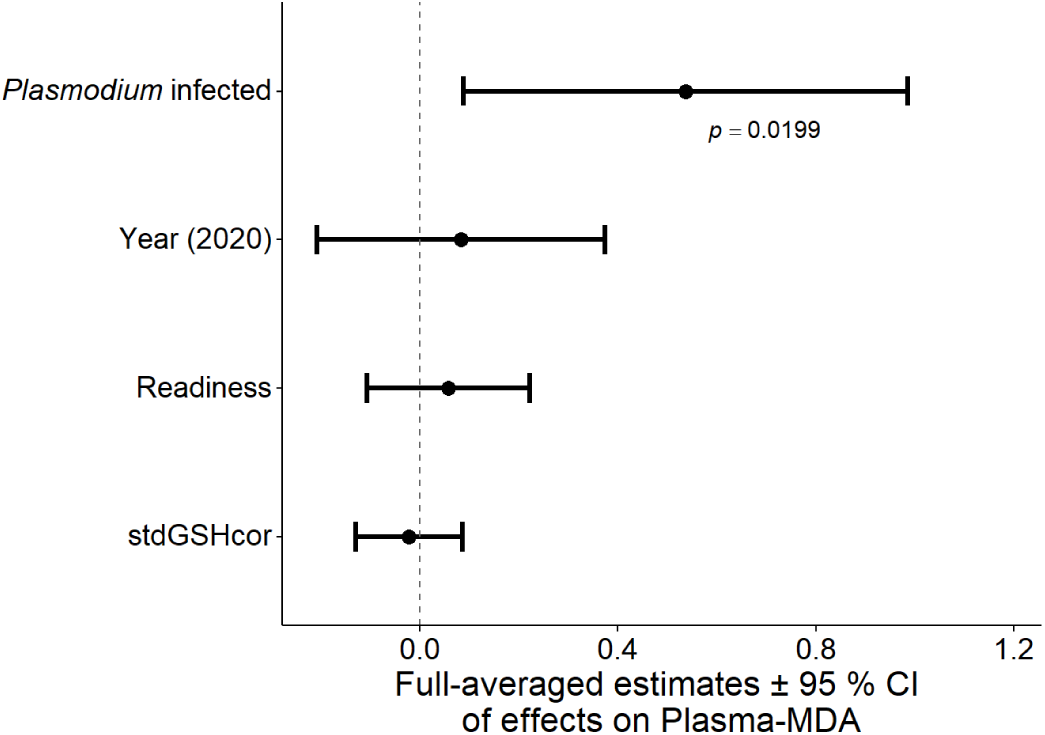
Standardized full model averaging estimates with 95 % confidence intervals of the effects of variables included in the best models for variation in plasma-MDA (log transformed) of blackcaps, when these were tested using *Plasmodium* infection as the parasite-related predictor (the null model was included among the best models when we used this variable to test parasite influences). The *p*-values of statistically significant effects are shown. Square root transformation was applied to tGSH.

## 3. Discussion

We found that young blackcaps during autumn migration were more often coinfected than adults, which reciprocally were more likely to harbour single parasites when they were infected. We also found that *Plasmodium* parasites were always present in coinfection with other parasites, something we would not have been able to detect without combining several techniques for parasite detection [50,60]. Globally, infected birds showed signs of poorer capacity to maintain oxidative balance during migration, as shown by lower levels of TAS in coinfected individuals compared to not infected ones, and higher oxidative damage to lipids in blackcaps infected by *Plasmodium* parasites. These effects, similar to the observed for young male blackcaps in an earlier study including part of our dataset [21], could be generalised to all blackcaps in this study as we did not find any effect of age or sex on the physiological status nor the body condition of migrating blackcaps. Nevertheless, given the persistent detection of *Plasmodium* parasites in co-infection, mainly in young blackcaps, and the potential for increased virulence in such cases [53,54,80,81], our findings support the idea that parasite infections during autumn migration have their strongest demographic impact during the first year of life of blackcaps.

Birds infected by *Plasmodium* parasites in our population had higher levels of damage to lipids in their plasma as measured by MDA values. Moreover, we found a gradual decrease in the antioxidant capacity of plasma with increasing number of parasites in blood [15], which led to a significant difference between not infected and coinfected individuals. The small size of this effect was probably due to high variability of TAS measurements among birds that are (1) analysed in an observational study and (2) undergoing profound changes of their oxidative balance associated with migration, which combines the physiological consequences of flight effort with metabolic and dietary adjustments during stopovers [21,26,82]. Taken together, these effects probably inflate the error term in our analyses, so that the observed effect and its progressivity from not infected to coinfected individuals may be evidence of a relevant influence of parasites on the oxidative balance of migrating blackcaps.

Our results suggest that multiple haemosporidian infections and *Plasmodium* infections could be especially harmful to blackcaps, as they might increase the cost of migration by impairing oxidative status. Other studies have also found an increased virulence of coinfections of avian haemosporidians [51,81], and our study contributes to unveil the physiological link between haemosporidian coinfection and impaired bird fitness. In blackcaps, we found that coinfections had similar effects on young and adult individuals, yet their population impact may be especially relevant during the first year of life.

Previous studies have found that coinfected individuals experience delayed development, which could make them less prepared for migration [54]. Our study shows that coinfections may also decrease their chances of successfully completing the first migration cycle and, consequently, joining the breeding population as recruits. Summed up, these selective filters may partly explain the lower prevalence of coinfections (which often included *Plasmodium* parasites) among adult blackcaps, although this observation could also be explained if adults had cleared most virulent infections acquired earlier in their life, and tolerate less aggressive parasites in chronic infections [83,84]. These two possible causes of age-related differences in the frequency of most virulent infections are not mutually exclusive.

In our study, we did not find any sexual differences neither in the infection probability nor in the physiological status of blackcaps, as reported in other study systems [46–48]. Birds are most exposed to the effects of sexual hormones during the breeding season, when these effects can unbalance their immune and physiological system making males more prone than females to infection [85] and oxidative stress [12,38]. However, circulating levels of sex hormones decrease outside the breeding season, which may attenuate the differences between males and females in physiological status and infection levels [32,86]. Such differences may be especially hard to detect during autumn migration if individuals harbour low intensity, chronic infections.

We found a syndrome of physiological variables (greater fat deposits, high haemoglobin level, high triglyceride content and lower uric acid level) indicative of blackcaps’ readiness for departure from the stopover site that has been previously described [74]. During stopovers, birds replenish their fat stores, stop using proteins to obtain energy (having less protein catabolic subproducts such as uric acid as a consequence [39,63]), and compensate the haemoglobin loss from the previous flight stage [62]. Contrary to what we expected, birds that were more physiologically ready for departure had also lower levels of antioxidant capacity of plasma. The total antioxidant capacity of plasma might decrease during stopover since in this period birds need to compensate the oxidative damage caused during flight through hyperphagia, which is linked to a lower metabolic rate to promote fattening [26]. Antioxidant capacity might also be lower at the end of the stopover because of the accumulation and usage of polyunsaturated fatty acids that are prone to oxidation [26]. Also, there was a positive correlation between tGSH levels and damage to lipids in RBC. This could be interpreted as a hormetic response to migration-induced oxidative stress [20,26], and it has been found in other species like European robins during migration [87]. In RBC, the lipid damage during stopover could be dealt with through the upregulation of endogenous antioxidants [88]. Therefore, blackcaps might be facing migration-induced oxidative stress that needs to be compensated during stopovers. When individuals migrate with infections, adaptive dietary shifts favouring antioxidant intake may be a mechanism to maintain oxidative balance, as we found in young males in our sample [21].

Our results suggest a combined effect of migration and haemosporidian parasites on the oxidative balance of both young and adult blackcaps. Previous studies also found individual differences in how haemosporidian parasites were associated with oxidative status of breeding birds [12,30]. Although, migration may be an important challenge, the link between haemosporidian infections and oxidative stress has been poorly studied during this period [28], and future longitudinal studies may be key to interpret the lifetime costs of carrying parasites during seasonal migration. While our data point towards a greater demographic impact of migration with parasites during the first year of life (when a large fraction of individuals migrates facing coinfections with harmful *Plasmodium* parasites), we could not uncover how parasites change their prevalence and impact as individuals age [22,40,89]. Moreover, in our study, most of the infections have probably already become chronic (3.30 parasites in 5000 RBC, SD = 4.33, in young males [21]), having fewer effects on the immune system and oxidative status of their hosts [84], but see [90]. From this perspective, parasites acquired during migration may have a stronger impact if the physiological cost of infection increases during acute parasitaemia [91], a cost which might promote the selection of stopover habitat where parasite impacts may be reduced. Supporting this idea, many species of migratory forest birds shift habitat preferences during migration, often selecting edge-dominated and early successional forests in autumn [92]. In these habitats, antioxidant-rich fleshy fruits are abundant [21,93,94], while parasite vectors tend to be scarce [95–98]. All in all, we are far from understanding the role of parasites in the ecophysiology of bird migration [28]. Here, we provide compelling evidence of oxidative costs of haemosporidian parasite infections in blackcaps during autumn migration. Similar costs may need to be considered during other life-history stages, as they may contribute to establish the physiological nexus between parasite infections and bird fitness.

## Acknowledgements

We thank the Ecophysiology lab from the National Museum of Natural Sciences for providing access to their facilities and to Lorenzo Pérez-Rodríguez for his advice with physiological analyses. This study was funded by grants CGL2017-82117-P and PID2020-116121GB-I00 funded by MCIN/AEI/ 10.13039/501100011033 and by “ERDF A way of making Europe” (to JPT). JLA was funded by a Juan de la Cierva-Formación 2018 (FJC2018-036465-I) granted by the Spanish Ministry of Science and Innovation. LJG was funded by a FPU grant (FPU17/04716) funded by the Spanish Ministry of Universities.

## Ethics statement

All handling and sampling procedures were authorised by Comunidad de Madrid through permits 10/060632.9/18, 10/058955.9/20 and 10/241922.9/21.

## Competing interests statement

The authors declare that they have no conflict of interest.

## Author contributions statement

Conceptualization: LJG, JPT, CR. Data Curation: LJG, JPT, CR. Formal Analysis: LJG, JLA, JPT, CR. Funding Acquisition: JPT. Investigation: LJG, JLA, JPT, CR. Methodology: LJG, JLA, JPT, CR. Project Administration: JPT, CR. Resources: JPT. Supervision: JPT, CR. Validation: CR, LJG. Writing – Original Draft: LJG. Writing – Review & Editing: LJG, JLA, JPT, CR.

## Data availability statement

Data available from the Dryad Repository: Pending DOI

## Notes

### Competing Interest Statement

The authors have declared no competing interest.

