## Supplementary material for "Blackcaps (*Sylvia atricapilla*) with blood parasites have impaired oxidative status during autumn migration"

### Supplementary Table S1

Table S1. Lineages of haemosporidian parasites from 130 infected blackcaps, distinguishing between adults and young individuals. Parasite lineages differ from one another by at least one base pair of the MalAvi barcode (478 bp of the Cytochrome *b* gene) and are assigned to taxonomic species when the MalAvi database provides this information [1]. Infections that could not be sequenced are included as "Unknown". The total number of parasite occurrences is higher than the number of infections due to coinfections. Twenty-four individuals were not infected. A minimum prevalence is calculated for each parasite identified based on the total number of individuals tested from each age (note that unknown parasites may belong to any of the lineages identified or to other lineages).

| Age | Genus | Lineage | Species | n | Prevalence |
|---|---|---|---|---|---|
| Adult | *Haemoproteus* | SYAT01 | *H. parabelopolskyi* | 12 | 23.52 % |
| | | SYAT02 | *H. parabelopolskyi* | 4 | 7.84 % |
| | | SYAT03 | *H. pallidulus* | 1 | 1.96 % |
| | | SYAT07 | *H. parabelopolskyi* | 13 | 25.49 % |
| | | SYAT13 | Undetermined | 1 | 1.96 % |
| | | Unknown *Haemoproteus* | Undetermined | 17 | |
| | *Plasmodium* | Unknown *Plasmodium* | Undetermined | 6 | |
| | *Leucocytozoon* | BT2 | Undetermined | 1 | 1.96 % |
| | | SFC8 | Undetermined | 3 | 5.88 % |
| | | SYAT22 | Undetermined | 1 | 1.96 % |
| | | Unknown *Leucocytozoon* | Undetermined | 1 | |
| Young | *Haemoproteus* | SYAT01 | *H. parabelopolskyi* | 38 | 36.89 % |
| | | SYAT02 | *H. parabelopolskyi* | 22 | 21.36 % |
| | | SYAT03 | *H. pallidulus* | 1 | 0.97 % |
| | | SYAT07 | *H. parabelopolskyi* | 12 | 11.65 % |
| | | SYAT10 | Undetermined | 1 | 0.97 % |
| | | SYAT13 | Undetermined | 6 | 5.83 % |
| | | WW2 | *H. majoris* | 1 | 0.97 % |
| | | Unknown *Haemoproteus* | Undetermined | 21 | |
| | *Plasmodium* | SYAT05 | *P. vaughani* | 2 | 1.94 % |
| | | Unknown *Plasmodium* | Undetermined | 14 | |
| | *Leucocytozoon* | BT2 | Undetermined | 2 | 1.94 % |
| | | SFC8 | Undetermined | 3 | 2.91 % |
| | | SYAT22 | Undetermined | 1 | 0.97 % |
| | | Unknown *Leucocytozoon* | Undetermined | 3 | |

### Supplementary Table S2

Table S2. Selection of the best models including the multiple infection status to explain variation in each physiological variable, according to the Akaike Information Criterion corrected by sample size (models with ΔAICc ≤ 2 compared to the best). When the null model was one of the best, we only report other models with lower AICc. Variables that were significant in the full averaged model are in bold.

| Best models for each parameter | df | AICc | ΔAICc | Weight |
|---|---|---|---|---|
| *Haemoglobin* | | | | |
| 1. year | 3 | 439.6 | 0.00 | 0.19 |
| 2. null model | 2 | 440.1 | 0.49 | 0.15 |
| *tGSH (sqrt)* | | | | |
| 1. **RBC-MDA** | 3 | 395.2 | 0.00 | 0.15 |
| 2. plasma-MDA (log) + **RBC-MDA** | 4 | 396.2 | 1.02 | 0.09 |
| 3. readiness + **RBC-MDA** | 4 | 396.7 | 1.44 | 0.07 |
| 4. age + **RBC-MDA** | 4 | 397.1 | 1.84 | 0.06 |
| 5. sex + **RBC-MDA** | 4 | 397.1 | 1.84 | 0.06 |
| *TAS (sqrt)* | | | | |
| 1. **coinfection** + **readiness** + **uric acid (sqrt)** | 6 | 251.1 | 0.00 | 0.06 |
| 2. **coinfection** + **readiness** + sex + **uric acid (sqrt)** | 7 | 251.3 | 0.20 | 0.05 |
| 3. **coinfection** + **readiness** + **uric acid (sqrt)** + year | 7 | 251.6 | 0.47 | 0.04 |
| 4. **coinfection** + **readiness** + sex + **uric acid (sqrt)** + year | 8 | 251.9 | 0.72 | 0.04 |
| 5. **coinfection** + **readiness** + RBC-MDA + **uric acid (sqrt)** | 7 | 252.3 | 1.19 | 0.03 |
| 6. **readiness** + **uric acid (sqrt)** | 4 | 252.5 | 1.32 | 0.03 |
| 7. **coinfection** + **readiness** + sex + RBC-MDA + **uric acid (sqrt)** | 8 | 252.5 | 1.36 | 0.03 |
| 8. **coinfection** + **readiness** + RBC-MDA + **uric acid (sqrt)** + year | 8 | 252.6 | 1.49 | 0.03 |
| 9. **readiness** + sex + **uric acid (sqrt)** | 5 | 252.7 | 1.56 | 0.03 |
| 10. **coinfection** + **readiness** + sex + RBC-MDA + **uric acid (sqrt)** + year | 9 | 252.8 | 1.71 | 0.02 |
| *Uric acid (sqrt)* | | | | |
| 1. coinfection | 4 | 431.0 | 0.00 | 0.18 |
| 2. null model | 2 | 431.6 | 0.64 | 0.13 |
| *Plasma-MDA (log)* | | | | |
| 1. readiness | 3 | 404.1 | 0.00 | 0.05 |
| 2. null model | 2 | 404.2 | 0.07 | 0.05 |
| *RBC-MDA* | | | | |
| 1. **tGSH (sqrt)** | 3 | 394.8 | 0.00 | 0.15 |
| 2. **tGSH (sqrt)** + year | 4 | 396.4 | 1.59 | 0.07 |
| *Triglycerides (log)* | | | | |
| 1. age | 3 | 421.2 | 0.00 | 0.17 |
| 2. age + year | 4 | 421.2 | 0.07 | 0.16 |
| 3. sex + year | 4 | 422.0 | 0.87 | 0.11 |

| | | | | |
|---|---|---|---|---|
| 4. age + sex | 4 | 422.0 | 0.88 | 0.10 |
| 5. age + sex + year | 5 | 422.1 | 0.93 | 0.10 |
| 6. year | 3 | 422.1 | 0.94 | 0.07 |
| 7. sex | 3 | 422.9 | 1.73 | 0.06 |
| 8. null model | 2 | 423.1 | 1.93 | 0.02 |

### Supplementary Table S3

Table S3. Selection of the best models including *Plasmodium* infections to explain variation in each physiological variable, according to the Akaike Information Criterion corrected by sample size (models with ΔAICc ≤ 2 compared to the best). When the null model was one of the best, we only report other models with lower AICc. Variables that were significant in the full averaged model are in bold.

| Best models for each parameter | df | AICc | ΔAICc | Weight |
|---|---|---|---|---|
| *Haemoglobin* | | | | |
| 1. year | 3 | 439.6 | 0,00 | 0.16 |
| 2. null model | 2 | 440.1 | 0.49 | 0.13 |
| *tGSH (sqrt)* | | | | |
| 1.**RBC-MDA** | 3 | 395.2 | 0.00 | 0.12 |
| 2. plasma-MDA (log) + **RBC-MDA** | 4 | 396.2 | 1.02 | 0.07 |
| 3. readiness + **RBC-MDA** | 4 | 396.7 | 1.44 | 0.06 |
| 4. age + **RBC-MDA** | 4 | 397.1 | 1.84 | 0.05 |
| 5. sex + **RBC-MDA** | 4 | 397.1 | 1.84 | 0.05 |
| *TAS (sqrt)* | | | | |
| 1. readiness + **uric acid (sqrt)** | 4 | 252.5 | 0.00 | 0.06 |
| 2. readiness + sex + **uric acid (sqrt)** | 5 | 252.7 | 0.24 | 0.05 |
| 3. readiness + **uric acid (sqrt)** + year | 5 | 253.5 | 1.03 | 0.03 |
| 4. *Plasmodium* + readiness + **uric acid (sqrt)** | 5 | 253.6 | 1.16 | 0.03 |
| 5. readiness + sex + **uric acid (sqrt)** + year | 6 | 253.8 | 1.32 | 0.03 |
| 6. readiness + RBC-MDA + **uric acid (sqrt)** | 5 | 253.9 | 1.43 | 0.03 |
| 7. **uric acid (sqrt)** | 3 | 254.1 | 1.60 | 0.03 |
| 8. readiness + sex + RBC-MDA + **uric acid (sqrt)** | 6 | 254.1 | 1.67 | 0.03 |
| 9. *Plasmodium* + readiness + sex + **uric acid (sqrt)** | 6 | 254.3 | 1.81 | 0.02 |
| 10. age + readiness + **uric acid (sqrt)** | 5 | 254.3 | 1.84 | 0.02 |
| *Uric acid (sqrt)* | | | | |
| 1. *Plasmodium* | 3 | 430.5 | 0.00 | 0.17 |
| 2. *Plasmodium* + sex | 4 | 430.7 | 0.19 | 0.16 |
| 3. null model | 2 | 431.6 | 1.05 | 0.10 |
| *Plasma-MDA (log)* | | | | |
| 1. ***Plasmodium*** + readiness | 4 | 400.5 | 0.00 | 0.07 |
| 2. ***Plasmodium*** | 3 | 400.7 | 0.20 | 0.06 |
| 3. ***Plasmodium*** + readiness + year | 5 | 401.1 | 0.59 | 0.05 |
| 4. ***Plasmodium*** + year | 4 | 401.1 | 0.63 | 0.05 |
| 5. ***Plasmodium*** + tGSH (sqrt) | 4 | 402.0 | 1.46 | 0.03 |
| 6. ***Plasmodium*** + readiness + tGSH (sqrt) | 5 | 402.0 | 1.47 | 0.03 |
| 7. ***Plasmodium*** + tGSH (sqrt) + year | 5 | 402.4 | 1.91 | 0.03 |
| *RBC-MDA* | | | | |
| 1. **tGSH (sqrt)** | 3 | 394.8 | 0.00 | 0.14 |

| | | | | |
|---|---|---|---|---|
| 2. **tGSH (sqrt)** + year | 4 | 396.4 | 1.59 | 0.06 |
| 3. *Plasmodium* + **tGSH (sqrt)** | 4 | 396.5 | 1.69 | 0.06 |
| *Triglycerides (log)* | | | | |
| 1. age | 3 | 421.2 | 0.00 | 0.13 |
| 2. age + year | 4 | 421.2 | 0.07 | 0.12 |
| 3. sex + year | 4 | 422.0 | 0.87 | 0.08 |
| 4. age + sex | 4 | 422.0 | 0.88 | 0.08 |
| 5. age + sex + year | 5 | 422.1 | 0.93 | 0.08 |
| 6. year | 3 | 422.1 | 0.94 | 0.08 |
| 7. age + *Plasmodium* | 4 | 422.3 | 1.12 | 0.07 |
| 8. age + *Plasmodium* + year | 5 | 422.5 | 1.33 | 0.06 |
| 9. sex | 3 | 422.9 | 1.73 | 0.05 |
| 10. null model | 2 | 423.1 | 1.93 | 0.05 |

### Supplementary Figure S1

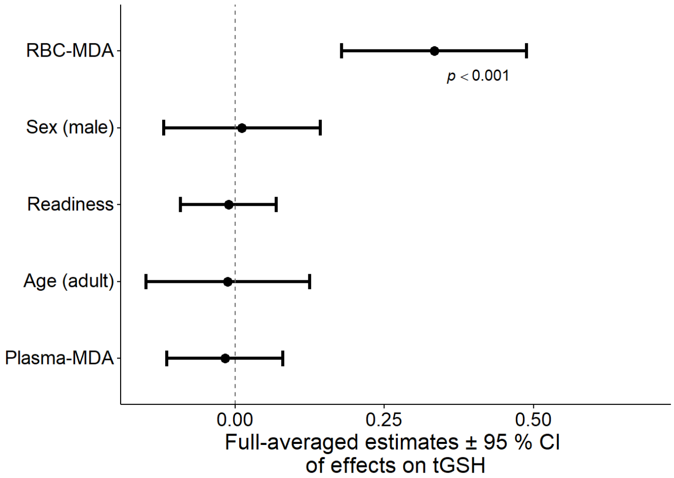


Figure S1. Standardized full model averaging estimates with 95 % confidence interval of the effects of variables included in the best models for variation in tGSH (square root transformed) of blackcaps. None of the best models included multiple infection status or *Plasmodium* infection as predictors. The *p*-values of statistically significant effects are shown. Plasma-MDA was log transformed.

### Supplementary Figure S2

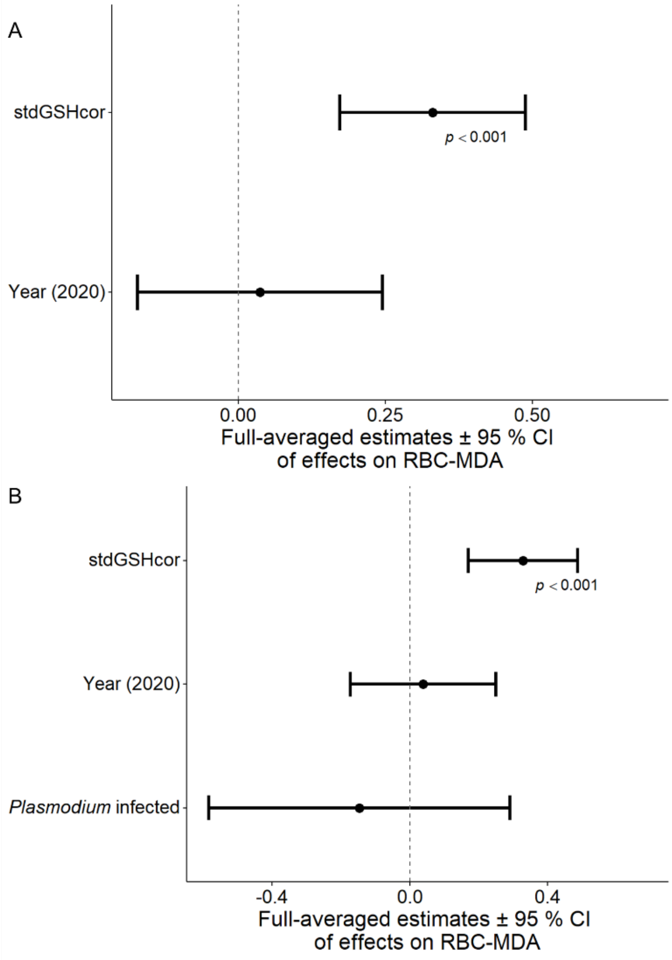


Figure S2. Standardized full model averaging estimates with 95 % confidence interval of the effects of variables included in the best models for variation in RBC-MDA of blackcaps, when these were tested using multiple infection status (A) or *Plasmodium* infection (B) as parasite-related predictors (none of the best models included multiple infection status as predictor when we tested the effect of parasites with this variable). The *p*-values of statistically significant effects are shown. The effect of multiple infection status is shown with not infected as the reference level. Square root transformation was applied to tGSH.
